# scDD: A statistical approach for identifying differential distributions in single-cell RNA-seq experiments

**DOI:** 10.1101/035501

**Authors:** Keegan D. Korthauer, Li-Fang Chu, Michael A. Newton, Yuan Li, James Thomson, Ron Stewart, Christina Kendziorski

## Abstract

The ability to quantify cellular heterogeneity is a major advantage of single-cell technologies. Although understanding such heterogeneity is of primary interest in a number of studies, for convenience, statistical methods often treat cellular heterogeneity as a nuisance factor. We present a novel method to characterize differences in expression in the presence of distinct expression states within and among biological conditions. Using simulated and case study data, we demonstrate that the modeling framework is able to detect differential expression patterns of interest under a wide range of settings. Compared to existing approaches, scDD has higher power to detect subtle differences in gene expression distributions that are more complex than a mean shift, and is able to characterize those differences. The freely available R package scDD implements the approach.

## Background

Coordinated gene expression is fundamental to an organism’s development and maintenance, and aberrations are common in disease. Consequently, experiments to measure expression on a genome-wide scale are pervasive. The most common experiment involves the quantification of mRNA transcript abundance averaged over a population of thousands or millions of cells. These so-called traditional, or bulk, RNA-seq experiments have proven useful in a multitude of studies. However, because bulk RNA-seq does not provide a measure of cell specific expression, many important signals go unobserved. A gene that appears to be expressed at a relatively constant level in a bulk RNA-seq experiment, for example, may actually be expressed in sub-groups of cells at levels that vary substantially (see Figure 1).

**Figure 1.**
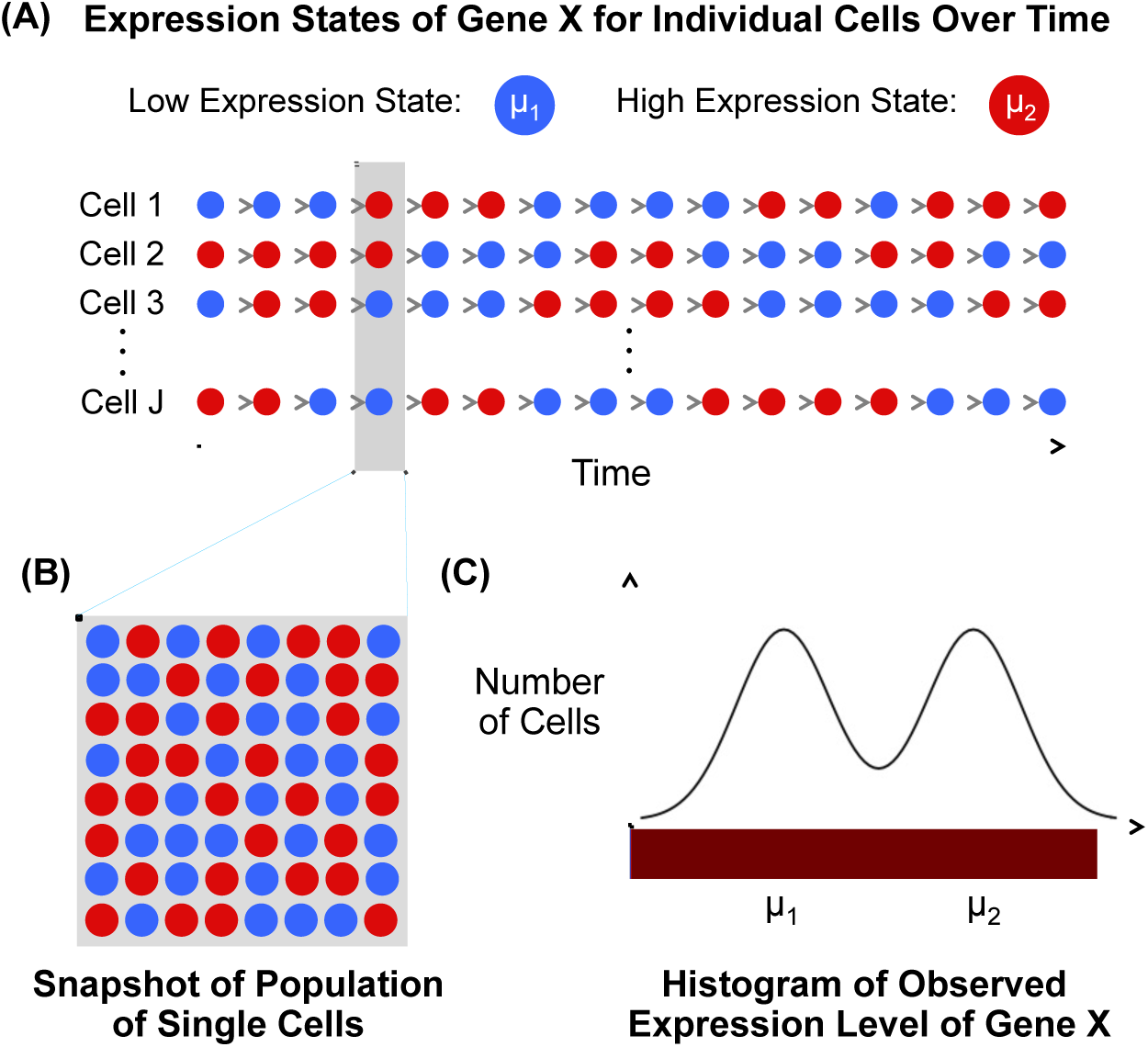
Schematic of the presence of two cell states within a cell population which can lead to bimodal expression distributions. (A) Time series of the underlying expression state of gene ☓ in a population of unsynchronized single cells, which switches back and forth between a low and high state with mean *μ*_1_ and *μ*_2_, respectively. The color of cells at each time point corresponds to the underlying expression state. (B) Population of individual cells shaded by expression state of gene ☓ at a snapshot in time. (C) Histogram of the observed expression level of gene ☓ for the cell population in (B).

Single-cell RNA-seq (scRNA-seq) facilitates the measurement of genome-wide mRNA abundance in individual cells, and as a result, provides the opportunity to study the extent of gene-specific expression heterogeneity within a biological condition, and the impact of changes across conditions. Doing so is required for discovering novel cell types [1, 2], for elucidating how gene expression changes contribute to development [3, 4, 5], for understanding the role of cell heterogeneity on the immune response [6, 7] and cancer progression [6, 8, 9, 10], and for predicting response to chemotherapeutic agents [11, 12, 13]. Unfortunately, the statistical methods available for characterizing gene-specific expression within a condition and for identifying differences across conditions in scRNA-seq are greatly limited, largely because they do not fully accommodate the cellular heterogeneity that is prevalent in single-cell data.

To identify genes with expression that varies across biological conditions in an scRNA-seq experiment, a number of early studies used methods from bulk RNA-seq [12, 10, 4, 14, 15]. In general, the methods assume that each gene has a latent level of expression within a biological condition, and that measurements fluctuate around that level due to biological and technical sources of variability. In other words, they assume that gene-specific expression is well characterized by a unimodal distribution within condition. Further, tests for differences in expression to identify so-called differentially expressed (DE) genes amount to tests for shifts in the unimodal distributions across conditions. A major drawback of these approaches in the single-cell setting is that, due to both biological and technical cell-to-cell variability, there is often an abundance of cells for which a given gene’s expression is unobserved [16, 7, 17] and, consequently, unimodal distributions are insufficient.

To address this, a number of statistical methods have been developed recently to accommodate bimodality in scRNA-seq data [17, 18]. In these mixture-model based approaches, one component distribution accommodates unobserved, or dropout, measurements (which include zero and, optionally, thresholded low-magnitude observations) and a second unimodal component describes gene expression in cells where expression is observed. Although these approaches provide an advance over unimodal models used in bulk, they are insufficient for characterizing multi-modal expression data, which is common in scRNA-seq experiments (see Figure 2).

**Figure 2.**
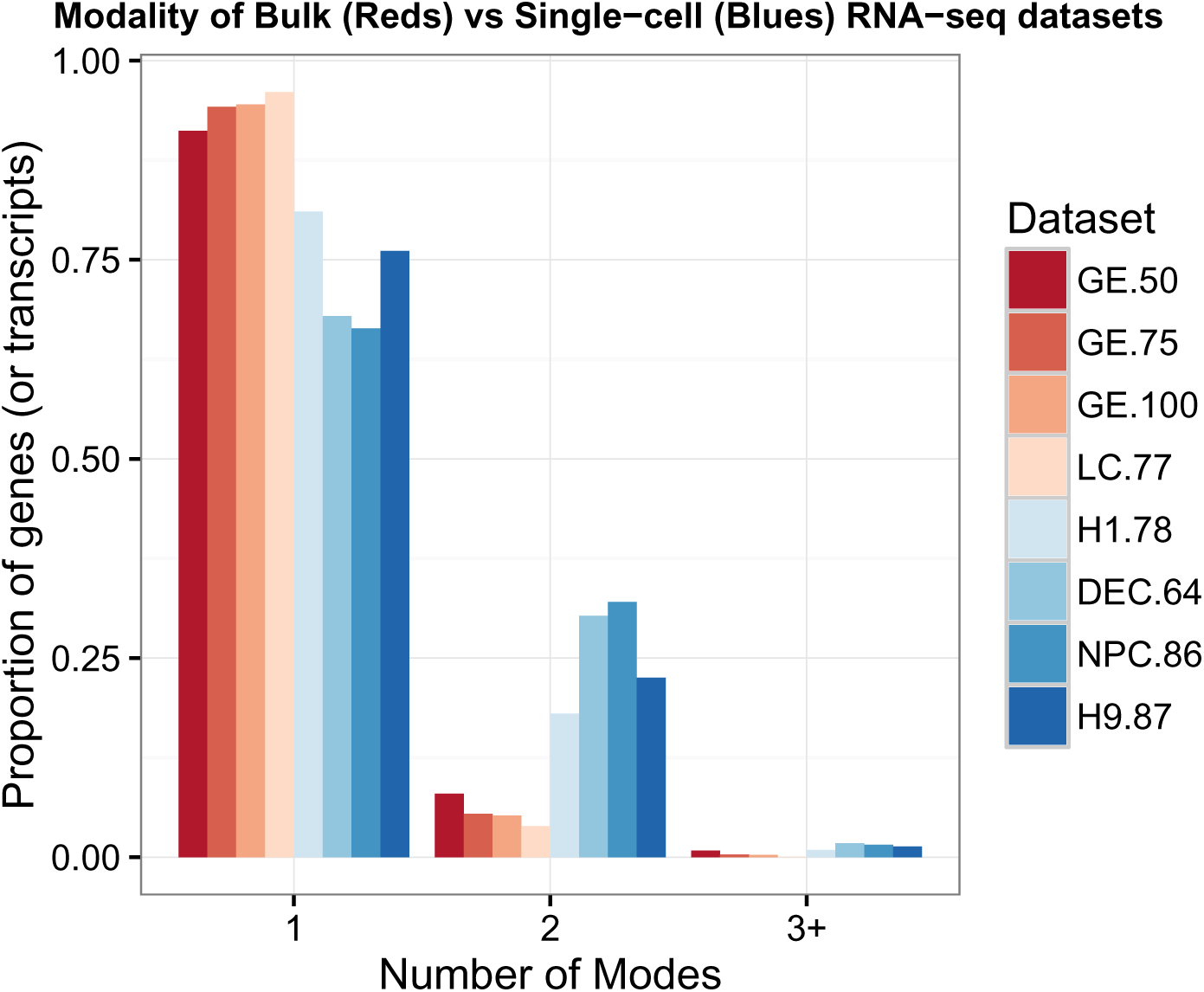
Bar plot of the proportion of genes (or transcripts) in each dataset where the log-transformed nonzero expression measurements are best fit by a 1, 2 or 3+ mode normal mixture model (where ‘3+’ denotes 3 or more). Modality is determined using a BIC selection criteria with filtering (see Partition estimation section). Red shades denote bulk RNA-seq datasets, and blue shades denote single-cell datasets. The number following each dataset label indicates the number of samples present (e.g. GE.50 is a bulk dataset with 50 samples). Datasets GE.50, GE.75, and GE.100 are constructed by randomly sampling 50, 75, and 100 samples from GEUVADIS [55]. Dataset LC consists of 77 normal samples from the TCGA lung adenocarcinoma study [56]. For details on the single-cell datasets, see Methods Section.

Specifically, a number of studies have shown that many types of heterogeneity can give rise to multiple expression modes within a given gene [19, 20, 21, 22, 23]. For example, there are often multiple states among expressed genes [19, 20, 22](a schematic is shown in Figure 1). The transition between cell states may be primarily stochastic in nature and result from expression bursts [24, 25], or result from positive feedback signals [26, 19, 23]. Beyond the existence of multiple stable states, multiple modes in the distribution of expression levels in a population of cells may also arise when the gene is either oscillatory and unsynchronized, or oscillatory with cellular heterogeneity in frequency, phase, and amplitude [21, 23].

Figure 3 illustrates common multi-modal distributions within and across biological conditions. When the overall mean expression level for a given gene is shifted across conditions, bulk methods, or recent methods for scRNA-seq [18, 17, 27, 28], may be able to identify the gene as showing some change. However, as we show here, they would be relatively underpowered to do so, and they would be unable to characterize the change, which is often of interest in an scRNA-seq experiment. For example, the gene in Figure 3 (C) shows a differential number of modes (DM), while the gene in Figure 3 (B) shows a differential proportion (DP) of cells at each expression level across conditions. Differentiating between DM and DP is important since the former suggests the presence of a distinct cell type in one condition, but not the other, while the latter suggests a change in splicing patterns among individual cells [7] or cell-specific responses to signaling [29].

**Figure 3.**
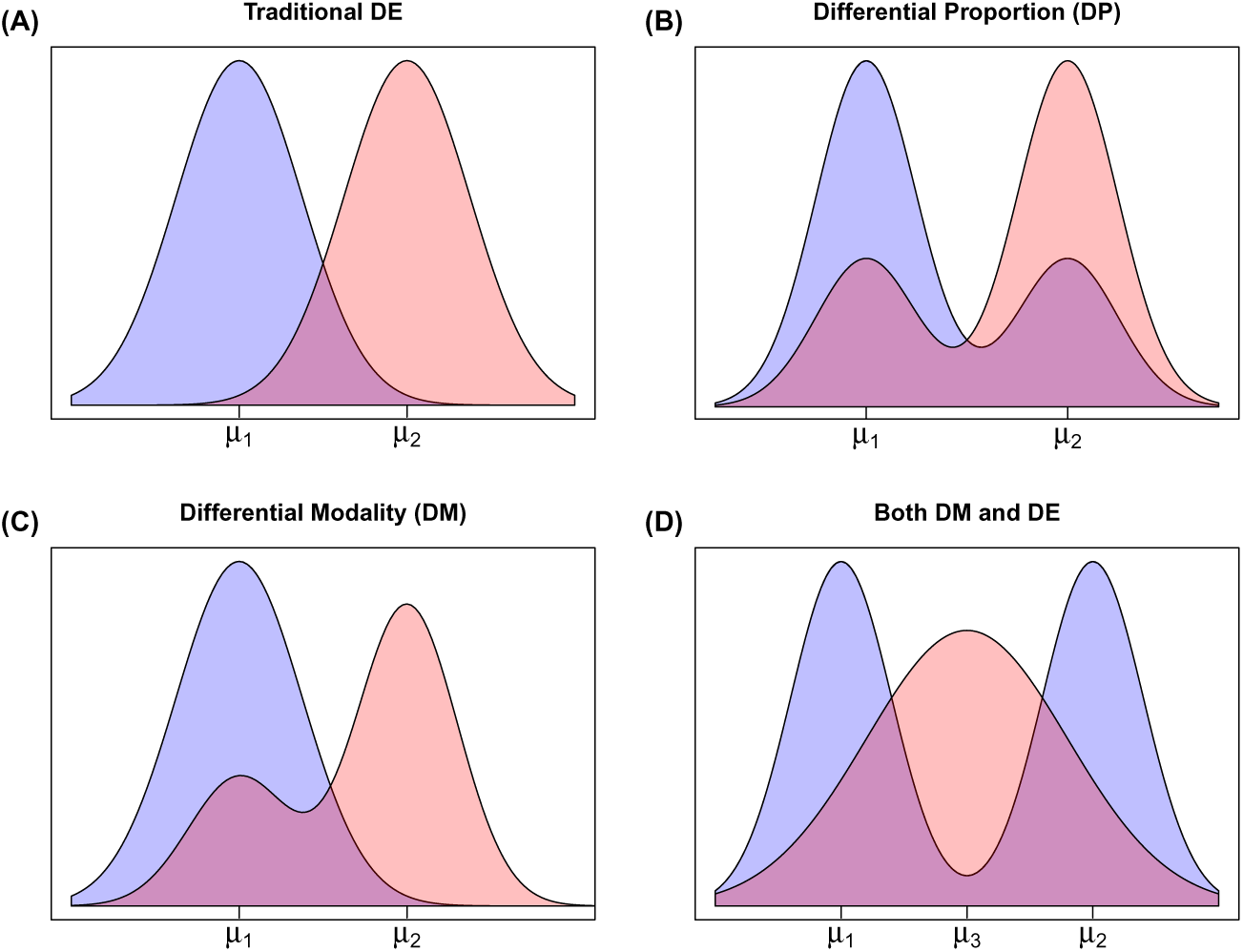
Diagram of plausible differential distribution patterns (histograms), including (A) traditional differential expression, (B) differential proportion within each mode, (C) differential modality, and (D) both differential modality and differential expression.

Here we develop a Bayesian modeling framework, scDD, to facilitate the characterization of expression within a biological condition, and to identify genes with differential distributions (DD) across conditions in an scRNA-seq experiment. A DD gene may be classified as DE, DM, DP, or both DM and DE (abbreviated DB; Figure 3 provides an overview of each pattern). Simulation studies suggest that the approach provides improved power and precision for identifying differentially distributed genes. Additional advantages are demonstrated in a case study of human embryonic stem cells.

## Results and discussion

### Human embryonic stem cell data

Single-cell RNA-seq data was generated in the James Thomson Lab at the Morgridge Institute for Research (see methods for details). Here we analyze data from two undifferentiated human embryonic stem cell (hESC) lines: the male H1 line (78 cells) and the female H9 line (87 cells). In addition, we include data from two differentiated cell types that are both derived from H1: definitive endoderm cell (DEC, 64 cells) and neuronal progenitor cell (NPC, 86 cells). The relationship between these four cell types is summarized by the diagram in Figure 4. As discussed in the case study results, it is of interest to characterize the differences in distributions of gene expression among these four cell types to gain insight into the genes that regulate the differentiation process.

**Figure 4.**
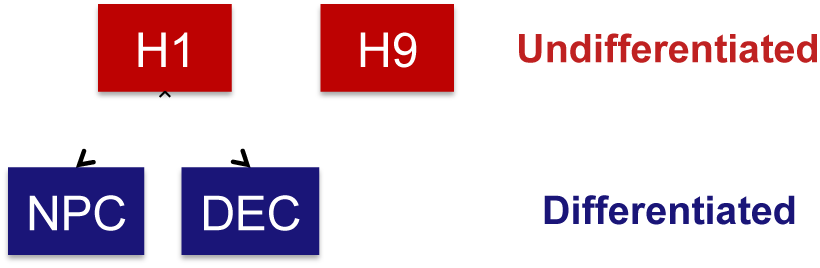
Relationship of cell types used in hESC case study. H1 and H9 are undifferentiated hESC lines. NPC (neuronal progenitor cells) and DEC (definitive endoderm cells) are differentiated cell types derived from HI.

### Publicly available human myoblast and mouse embryonic stem cell data

We also apply our method to two publicly available scRNA-seq datasets to determine which genes are differentially distributed following stimulation or inhibition of differentiation via specialized growth medium. Using data from [30], we compare gene expression of human myoblast cells cultured in standard growth medium (T0, 96 cells) with those treated with differentiation-inducing medium for 72 hours (T72, 84 cells). Additionally, we use data from [31] to compare gene expression of mouse embryonic stem cells (mESC) cultured in standard medium (Serum+LIF, 93 cells) with those cultured on differentiation-inhibiting medium (2i+LIF, 94 cells).

### Simulated data

We evaluate model performance using log-transformed count data simulated from mixtures of negative binomial distributions. The analysis of log-transformed counts from bulk RNA-seq has been shown to perform as well as utilizing count-based modeling assumptions [32, 33]. Recent scRNA-seq analyses have also assumed normality of log-transformed nonzero measurements [7, 18]. For each simulated dataset, 10,000 genes were simulated for two conditions with four different sample size settings (50, 75, 100, and 500 cells in each condition). The majority of the genes (8,000) were simulated out of the same model in each condition, and the other 2,000 represent genes with the four types of differential distributions (DD) outlined in Figure 3. The 2,000 DD genes were split equally into the following four categories:

- DE: single component with different mean in each condition
- DP: two components in each condition with equal component means across conditions; the proportion in the low mode is 0.33 for condition 1 and 0.66 for condition 2
- DM: single component in condition 1; two components in condition 2 with one overlapping component. Half of the condition 2 cells belong to each mode
- DB: single component in condition 1; two components in condition 2 with no overlapping components. The mean of condition 1 is half-way between the means in condition 2. Half of the cells in condition 2 belong to each mode.

where different clusters (with different generating distributions) are referred to as components, and different biological groups of interest are referred to as conditions.

Of the 8,000 null genes, 4,000 were generated from a single negative binomial component (EE) and the other 4,000 from a two-component binomial mixture (EP). Parameters of the negative binomial distributions for the unimodal genes were chosen to be representative of the observed means and variances in the H1 dataset. Fold-changes for DE genes were chosen to be representative of those observed in the H1 and DEC comparison. Distances between (log-scale) component means Δ_*μ*_σ̂ in the multi-modal genes were varied for the two-component cases, with equal proportion of genes at each setting of Δ_*μ*_ ∈ {2, 3, 4, 5, 6}, where σ̂ is the estimated cluster-specific standard deviation. More details are provided in the Methods section.

## The scDD modeling framework

Let *Y*_*g*_ = (*y*_*g*1_,…,*y*_*g*_*J*) be the log-transformed nonzero expression measurements of gene *g* in a collection of *J* cells from two biological conditions. We assume that measurements have been normalized to adjust for technical sources of variation including amplification bias and sequencing depth. Under the null hypothesis of equivalent distributions (i.e. no dependence on condition), we let *Y*_*g*_ be modeled by a conjugate Dirichlet process mixture (DPM) of normals (see Methods section for more details). Gene *g* may also have expression measurements of zero in some cells; these are modeled as a separate distributional component (see section ‘Differential proportion of zeroes’ for more details).

Ultimately, we would like to calculate a Bayes Factor for the evidence that the data arises from two independent condition-specific models (differential distributions (DD)) versus one overall model that ignores condition (equivalent distributions (ED)). Let ***M***_*DD*_ denote the differential distributions hypothesis, and ***M***_*ED*_ denote the equivalent distributions hypothesis. A Bayes Factor in this context for gene *g* would be:

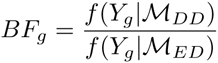

where *f* (*Y*_*g*_|***M***) denotes the predictive distribution of the observations from gene *g* under the given hypothesis. In general, there is no analytical solution for this distribution under the Dirichlet process mixture model framework. However, under the Product Partition Model (PPM) formulation (see Methods section for more details), we can get a closed form solution for *f* (*Y*_*g*_, *Z*_*g*_|***M***), where *Z*_*g*_ represents a partition of samples to mixture components. As the partition *Z*_*g*_ cannot be integrated out, we introduce an approximate Bayes Factor score:

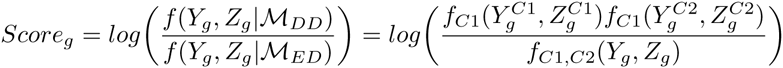

where *C*1 and *C*2 denote condition 1 and 2, respectively, and the score is evaluated at the partition estimate *Ẑ*_*g*_. A high value of this score presents evidence that a given gene is differentially distributed. Significance of the score is assessed via a permutation test. Specifically, condition labels are permuted and partition estimates are obtained within the new ‘conditions’. For each permuted data set, the Bayes Factor score is calculated; the default in scDD is 1,000 permutations. For each gene, an empirical p-value is calculated, and FDR is controlled for a given target value using the method of [34].

If covariates are available, instead of permuting the observed values, the relationship between the clustering and covariates can be preserved by permuting the residuals of a linear model that includes the covariate and using the fitted values [35]. As pointed out by [18], the cellular detection rate is a potential confounder variable, so the permutation procedure in the case studies is adjusted in this manner. If other known confounders exist and are measured, these can also be incorporated in the same manner.

### Classification of significant DD genes

For genes that are identified as DD by the Bayes Factor score, of interest is classifying them into four categories that represent the distinct DD patterns shown in Figure 3. To classify the DD genes into these patterns (DE, DM, DP, and DB), scDD utilizes the conditional posterior distribution of the cluster-specific mean parameters given in Equation 6 (see Methods section). Posterior sampling is carried out to investigate the overlap of clusters across conditions. Let *c*_1_ be the number of components in condition 1, *c*_2_ the number of components in condition 2, and *c*_*OA*_ the number of components overall (when pooling condition 1 and 2). Only components containing at least 3 cells are considered in order to minimize the impact of outlier cells. Note that for interpretability, a DD gene must satisfy: *c*_1_ + *c*_2_ ≥ *c*_*OA*_ ≥ *min*(*c*_1_, *c*_2_). These bounds on the number of components overall represent the two extreme cases: condition 1 does not overlap with condition 2 at all, versus one condition completely overlaps with the other. Any cases outside of these boundaries are not readily interpretable in this context. The actions to take for all other possible combinations of *c*_1_, *c*_2_, and *c*_*OA*_ are detailed in the Methods section.

### Differential proportion of zeroes

For those genes that do not show differential distributions in the nonzero values, scDD allows a user to evaluate whether the proportion of zeroes differs significantly between the two conditions. This evaluation is carried out using logistic regression adjusted for the proportion of genes detected in each cell as in [18]. Genes with a *χ*^*2*^ test p-value of less than 0.025 (after adjustment for multiple comparisons using the method of [34]) are considered to have a differential proportion of zeroes (DZ).

### Simulation study

A simulation study was conducted to assess the performance of scDD to identify DD genes, and to classify them as DE, DP, DM, or DB. Model performance on the simulated data was assessed based on (1) the ability to estimate the correct number of components, (2) the ability to detect significantly DD genes, and (3) the ability to classify DD genes into their correct categories. These three criteria are explored in the next three sections, respectively. Existing methods for differential expression analysis are also evaluated for the second criteria.

#### Estimation of the number of components

We first examine the ability of scDD to detect the correct number of components. Table 1 displays the proportion of bimodal and unimodal simulated genes where the correct number of components was identified. For bimodal genes, results are stratified by cluster mean distance. It is clear that the ability of the algorithm to correctly identify the correct number of components in bimodal genes improves as the component mean distance or sample size increases. The results for unimodal genes are not as sensitive to sample size, however the proportion of genes identified as bimodal increases slightly with more samples. We conclude that the partition estimate is able to reliably detect the true number of components for reasonable sample and effect sizes.

**Table 1.** Rate of detection of correct number of components in simulated data

| Sample<br>Size | Bimodal |  |  |  |  | Unimodal |
| --- | --- | --- | --- | --- | --- | --- |
| | Cluster mean distance $\Delta_\mu$ | | | | | |
|  | 2 | 3 | 4 | 5 | 6 |  |
| 50 | 0.056 | 0.196 | 0.579 | 0.848 | 0.922 | 0.907 |
| 75 | 0.052 | 0.252 | 0.719 | 0.917 | 0.957 | 0.908 |
| 100 | 0.050 | 0.302 | 0.811 | 0.950 | 0.974 | 0.905 |
| 500 | 0.073 | 0.417 | 0.959 | 0.995 | 0.991 | 0.884 |

Average proportion of simulated bimodal and unimodal genes where the correct number of components was identified, averaged over gene category and condition. Averages are calculated over 20 replications. Standard errors were $< 0.025$ (not shown).

#### Detection of DD genes

Next, we examine the ability of scDD to identify the non-null genes as significantly DD, and compare to the existing methods SCDE [17] and MAST [18]. For each method, target FDR was set at 5% (see Methods for details). The power to detect each gene pattern as DD for all three methods is shown in Table 2. Note that the calculations here are taken before the classification step for scDD, so power is defined as the proportion of genes from each simulated category that are detected as DD. In general, the power to detect DD genes improves with increased sample size for all three methods. Our approach has comparable power to SCDE and MAST for DE and DM genes, but higher overall power to detect DP and DM genes. Interestingly, SCDE has very low power to detect DP genes, whereas MAST shows very low power to detect DB genes.

**Table 2.** Power to detect DD genes in simulated data

| Sample Size | Method | True Gene Category |  |  |  | Overall (FDR) |
| --- | --- | --- | --- | --- | --- | --- |
|  |  | DE | DP | DM | DB |  |
| 50 | scDD | 0.893 | **0.418** | **0.898** | **0.572** | **0.695** (0.029) |
|  | SCDE | 0.872 | 0.026 | 0.817 | 0.260 | 0.494 (0.004) |
|  | MAST | **0.908** | 0.400 | 0.871 | 0.019 | 0.550 (0.026) |
| 75 | scDD | 0.951 | 0.590 | **0.960** | **0.668** | **0.792** (0.031) |
|  | SCDE | 0.948 | 0.070 | 0.903 | 0.387 | 0.577 (0.003) |
|  | MAST | **0.956** | **0.633** | 0.943 | 0.036 | 0.642 (0.022) |
| 100 | scDD | 0.972 | 0.717 | **0.982** | **0.727** | **0.850** (0.033) |
|  | SCDE | 0.975 | 0.125 | 0.946 | 0.478 | 0.631 (0.003) |
|  | MAST | **0.977** | **0.752** | 0.970 | 0.045 | 0.686 (0.022) |
| 500 | scDD | **1.000** | 0.983 | **1.000** | **0.905** | **0.972** (0.035) |
|  | SCDE | **1.000** | 0.855 | 0.998 | 0.787 | 0.910 (0.004) |
|  | MAST | **1.000** | **0.993** | **1.000** | 0.170 | 0.791 (0.022) |

Average power to detect simulated DD genes by true category. Averages are calculated over 20 replications. Standard errors were $< 0.025$ (not shown).

#### Classification of DD genes

Next, we examine the ability of scDD to classify each DD gene into its corresponding category. Table 3 shows the Correct Classification Rate in each category for DD genes that were correctly identified during the detection step (calculated as the proportion of true positive genes detected as DD for a given category that were classified into the correct category). The classification rates do not depend strongly on sample size, with the exception of DP, which decreases with increasing sample size. This decrease results from an increase in the DD detection rate of DP genes with small cluster mean distance, which have a lower correct classification rate (as shown below).

**Table 3.** Correct Classification Rate in simulated data

| Sample Size | Gene Category |  |  |  |
| --- | --- | --- | --- | --- |
|  | DE | DP | DM | DB |
| 50 | 0.719 | 0.801 | 0.557 | 0.665 |
| 75 | 0.760 | 0.732 | 0.576 | 0.698 |
| 100 | 0.782 | 0.678 | 0.599 | 0.706 |
| 500 | 0.816 | 0.550 | 0.583 | 0.646 |

Average Correct Classification Rate for detected DD genes. Averages are calculated over 20 replications. Standard errors were $< 0.025$ (not shown).

Since the ability to correctly classify a DD gene depends on the ability to detect the correct number of components (see classification algorithm in Methods), we also examine how the Correct Classification Rate varies with cluster mean distance for the categories that contain bimodal genes (DP, DM, and DB). As shown in Table 4, the classification rates improve as Δ_*μ*_ increases. This pattern mirrors the trend in Table 1, and suggests that misclassification events occur largely due to incorrect estimation of the number of components. Performance generally increases with sample size, especially at lower values of Δ_*μ*_. In general, the ability of the algorithm to classify detected DD genes into their true category is robust when components are well-separated and improves with increasing sample size.

**Table 4.** Average correct classification rates by cluster mean distance

| Sample Size | Gene Category | Cluster mean distance $\Delta_\mu$ | | | | |
| --- | --- | --- | --- | --- | --- | --- |
|  |  | 2 | 3 | 4 | 5 | 6 |
| 50 | DP | 0.02 | 0.20 | 0.78 | 0.94 | 0.98 |
|  | DM | 0.10 | 0.23 | 0.59 | 0.81 | 0.89 |
|  | DB | 0.08 | 0.22 | 0.59 | 0.80 | 0.80 |
| 75 | DP | 0.02 | 0.18 | 0.77 | 0.94 | 0.97 |
|  | DM | 0.08 | 0.27 | 0.69 | 0.86 | 0.90 |
|  | DB | 0.09 | 0.29 | 0.71 | 0.83 | 0.84 |
| 100 | DP | 0.03 | 0.16 | 0.74 | 0.93 | 0.95 |
|  | DM | 0.10 | 0.32 | 0.76 | 0.87 | 0.91 |
|  | DB | 0.08 | 0.32 | 0.80 | 0.85 | 0.84 |
| 500 | DP | 0.01 | 0.15 | 0.72 | 0.91 | 0.93 |
|  | DM | 0.12 | 0.33 | 0.72 | 0.85 | 0.89 |
|  | DB | 0.03 | 0.43 | 0.85 | 0.85 | 0.85 |

Average Correct Classification Rates stratified by $\Delta_\mu$ . Averages are calculated over 20 replications. Standard errors were $< 0.025$ (not shown).

### Case study: identifying DD genes between hESC types

The comprehensive characterization of transcriptional dynamics across hESC lines and derived cell types aims to provide insight into the gene regulatory processes governing pluripotency and differentiation [36, 37, 38]. Previous work utilizing microarrays and bulk-RNA sequencing largely focused on identifying genes with changes in average expression level across a population of cells. By examining transcriptional changes at the single cell level, we can uncover global changes that were undetectable when averaging over the population. In addition, we gain the ability to assess the level of heterogeneity of key differentiation regulators, which may lead to the ability to assess variation in pluripotency [39] or differentiation potential of individual cells.

The number of significant DD genes for each cell type comparison is shown in Table 5 for scDD, SCDE, and MAST. Note that the comparison of H1 and H9 detects the fewest number of DD genes for all three methods, a finding that is consistent with the fact that both of these are undifferentiated hESC lines and it is expected that they are the most similar among the comparisons. In all four comparisons, the number of genes identified by our method is greater than SCDE and similar to MAST.

**Table 5.** Number of DD genes identified in the hESC case study data for scDD, SCDE, and MAST. Note that the Total for scDD includes genes detected as DD but not categorized.

| Comparison | scDD |  |  |  |  |  | SCDE | MAST |
| --- | --- | --- | --- | --- | --- | --- | --- | --- |
|  | DE | DP | DM | DB | DZ | Total |  |  |
| H1 vs NPC | 1686 | 270 | 902 | 440 | 1603 | 5555 | 2921 | 5887 |
| H1 vs DEC | 913 | 254 | 890 | 516 | 911 | 5295 | 1616 | 3724 |
| NPC vs DEC | 1242 | 327 | 910 | 389 | 2021 | 5982 | 2147 | 5624 |
| H1 vs H9 | 260 | 55 | 85 | 37 | 145 | 739 | 111 | 1119 |

Figure 5 (A) displays top-ranked genes for each category that are not identified by MAST or SCDE for the H1 versus DEC comparison. Among the genes identified exclusively by scDD for the H1 versus DEC comparison are *CHEK2,* a cell-cycle checkpoint kinase [40], and *CDK7,* a cyclin-dependent kinase that plays a key role in cell cycle regulation through the activation of other cyclin-dependent kinases [41]. It has been shown that embryonic stem cells express cyclin genes constitutively, whereas in differentiated cells cyclin levels are oscillatory [42]. This finding is consistent with the differential modality of the *CDK7* gene shown in Figure 5 (B). Similarly, scDD identifies several genes involved in the regulation of pluripotency that are not identified by the other two methods (Figure 5 (C)). For example, *FOXP1* exhibits alternative splicing activity in hESCs, stimulating expression of several key regulators of pluripotency [43]. The *PSMD12* gene encodes a subunit of the proteasome complex which is vital to maintenance of pluripotency and has shown decreased expression in differentiating hESCs [44]. Both of these genes are also differentially distributed between H1 and the other differentiated cell type NPC.

**Figure 5.**
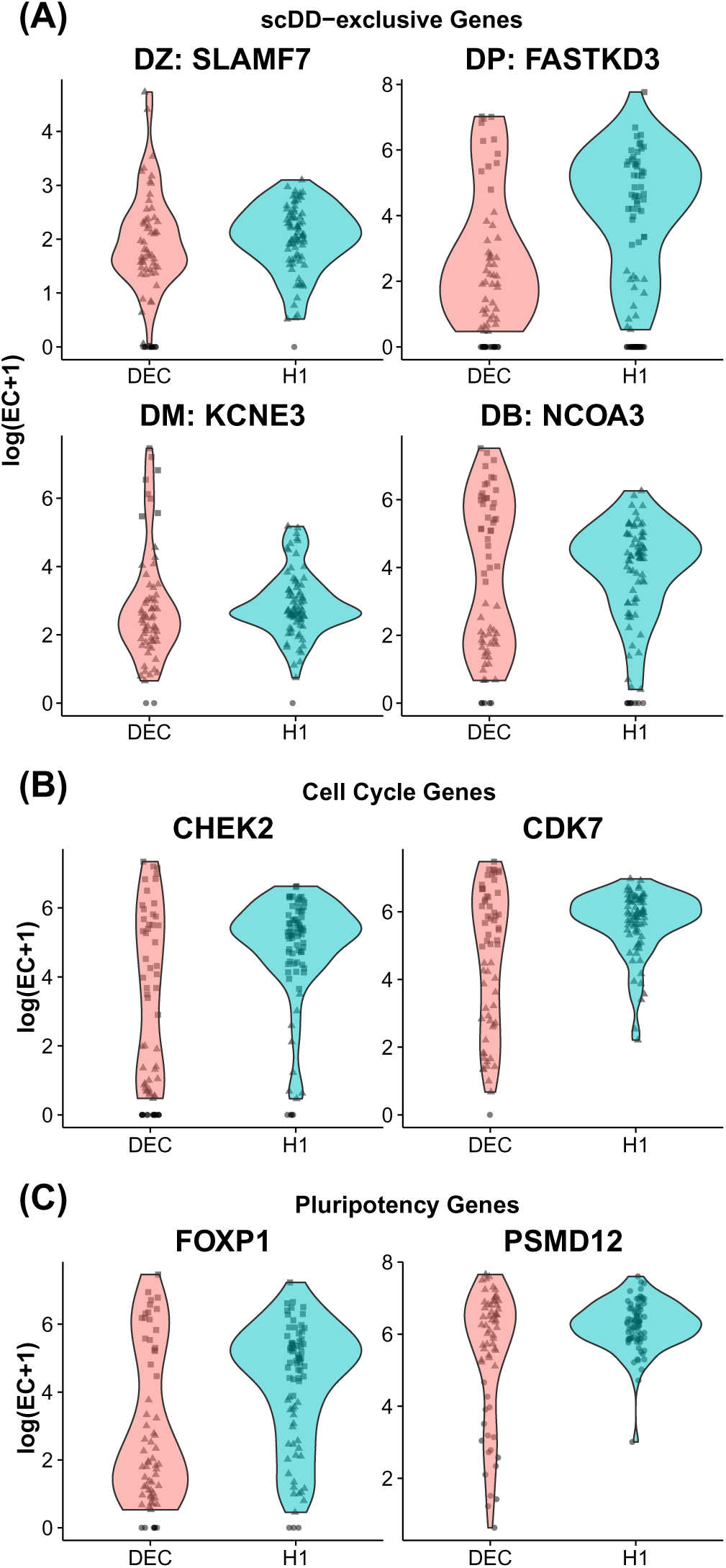
Violin plots (smoothed non-parametric kernel density estimates) for DD genes identified between H1 and DEC. Individual observations are displayed with jitter. Within a condition, points with the same shape are predicted to belong to the same cluster. (A) scDD-exclusive genes: representative genes from each category (DZ, DP, DM, DB) that are not detected by MAST or SCDE. Selected genes are top-ranked by permutation p-value in each category (DP, DM, DB) or had a significant χ^2^ test for a difference in the proportion of zeroes (DZ). (B) Cell Cycle genes: DD genes involved in cell cycle regulation (not detected by MAST or SCDE). (C) Pluripotency genes: DD genes involved in pluripotency regulation (not identified by MAST or SCDE).

In general, the vast majority of the genes found exclusively by scDD are categorized as something other than DE (ranging from 98.3% to 100% in the three case studies, see Supplementary Table S3), which suggests that they are predominantly characterized by differences that are more complex than the traditional DE pattern. The genes identified by MAST but not scDD are overwhelmingly characterized as those with a weak signal in both the nonzero and zero components (see Supplementary Figure S9), which can be difficult to interpret (see Supplement Section 3 for more details).

### Additional case studies

We also applied scDD and MAST to two additional case studies (the number of significant DD genes for each comparison are displayed in Table 6). SCDE was not used to analyze these datasets since it is intended for use on raw count data and the processed data made available by the authors of [30] and [31] were already normalized by FPKM and TPM, respectively. Similar to the results of the hESC case study, MAST and scDD identify similar numbers of significant genes. The genes that scDD finds exclusively are predominantly characterized by something other than a mean shift, a result which is also consistent with the hESC case study (see Supplementary Table S4).

**Table 6.** Number of DD genes identified in the myoblast and mESC case studies for scDD and MAST. Note that the Total for scDD includes genes detected as DD but not categorized.

| Comparison | scDD |  |  |  |  |  | MAST |
| --- | --- | --- | --- | --- | --- | --- | --- |
|  | DE | DP | DM | DB | DZ | Total |  |
| Myoblast: T0 vs T72 | 312 | 44 | 200 | 36 | 1311 | 2134 | 2904 |
| mESC: Serum vs 2i | 5233 | 76 | 1259 | 1128 | 670 | 9130 | 9706 |

### Advantages and limitations of the approach

We stress that our approach is inherently different from a method that detects traditional differential expression, such as [17] and [18] which aim to detect a shift in the mean of the expressed values. In addition to identifying genes that have differential distributions across conditions, our modeling framework allows us to identify subpopulations within each condition that have differing levels of expression of a given gene (i.e. which cells belong to which component). For such genes, the clustering automatically provides an estimate of the proportion of cells in each condition that belong to each subpopulation. We also do not require specification of the total number of components, which can vary for each gene.

When applied to cells at different differentiation stages, this information may provide insight into which genes are responsible for driving phenotypic changes. The gene in Figure 3(B), for example, shows a differential proportion (DP) of cells across conditions which is important to recognize since DP suggests a change in cell-specific responses to signaling [29, 7]. This is in contrast to the differential modes (DM) gene in Figure 3(C), which indicates the presence of a distinct cell type in one condition, but not the other. Recent methods for scRNA-seq [45, 18, 17, 27, 28] may be able to identify genes such as those shown in Figure 3(B-D) as differing between conditions. However, our simulations suggest that they would be relatively underpowered to do so, and they would be unable to characterize the change as DP, DM, or DB.

We also show through simulation that our approach can accommodate large sample sizes of several hundreds of cells per condition. Note, however, that the real strength in the modeling framework lies in the ability to characterize patterns of differential distributions. In the presence of extreme sparsity, this will be a challenge, since the number of nonzero observations in a given gene will be small. If the sample size of nonzero measurements is too small, it will be difficult to infer the presence of multiple underlying cell states. In practice, for larger and more sparse datasets it is recommended to verify that the number of cells expressing a given gene is in the range of the sample sizes considered in this study in order to fully utilize the available features of scDD.

The approach is limited in that adjustments for covariates are not directly incorporated into the model. In general, when the relationship between a potential confounding variable and the quantification of expression is well-known (e.g. increased sequencing depth is generally associated with increased expression measurements), this should be accounted for in a normalization procedure. For other covariates that are not as well-characterized (e.g. cellular detection rate, batch effects), residuals can be used in the permutation procedure, though a more unified approach would be desirable. Additionally, the approach is limited in that only pairwise comparisons across biological conditions are feasible. While an extended Bayes Factor score to test for dependence of condition on clustering for more than two conditions would be straightforward, the classification into meaningful patterns would be less so, and work is underway in that direction.

## Conclusions

To our knowledge, we have presented the first statistical method to detect differences in scRNA-seq experiments that explicitly accounts for potential multi-modality of the distribution of expressed cells in each condition. Such multi-modal expression patterns are pervasive in scRNA-seq data and are of great interest since they represent biological heterogeneity within otherwise homogeneous cell populations; and differences across conditions imply differential regulation or response in the two groups. We have introduced a set of five interesting patterns to summarize the key features that can differ between two conditions. Using simulation studies, we have shown that our method has comparable performance to existing methods when differences (mean shifts) exist between unimodal distributions across conditions, and outperforms existing approaches when there are more complex differences.

## Methods

### Software implementations and applications

All analyses were carried out using R version 3.1.1 [46]. The method MAST [18] was implemented using the MAST R package version 0.931, obtained from Github at https://github.com/RGLab/MAST. The adjustment for cellular detection rate as recommended in [18] was included in the case study, but not in the simulation study (only the ‘normal’ component of the test was considered here since no difference in dropout rate was simulated). The method SCDE [17] was implemented using the scde R package version 1.0, obtained from http://pklab.med.harvard.edu/scde/index.html. Since SCDE requires raw integer counts as input, and expected counts are non-integer valued, the ceiling function was applied to the unnormalized counts. For each approach, target FDR was controlled at 5%. Specifically, both MAST and SCDE provide gene-specific p-values and use the method of [34] to control FDR. We followed the same procedure here.

Our method is implemented using version 1.1.0 of the scDD R package, available at https://github.com/kdkorthauer/scDD. The analysis involves a computationally intensive permutation step which is executed in parallel on multiple cores if available. On a linux machine using 12 cores and up to 16 gigabytes of memory, this step took approximately 60 minutes for 1000 permutations of 1000 genes in the simulation of 50 samples per condition. Computation time scales approximately linearly with sample size, where this same task takes approximately 90 minutes for 100 samples per condition, and 300 minutes for sample size 500 per condition.

### hESC culture and differentiation

All cell culture and scRNA-seq experiments were conducted as described previously [47]. Briefly, undifferentiated H1 and H9 human ES cells were routinely maintained at the undifferentiated state in E8 medium on Matrigel (BD Bioscience) coated tissue culture plates with daily medium feeding [48]. Human ES cells were passaged every 3 to 4 days with 0.5 mM EDTA in PBS at 1:10 to 1:15 ratio for maintenance. H1 were differentiated according to previously established protocols [49, 50]. All the cell cultures performed in our laboratory have been routinely tested as negative for mycoplasma contamination.

For DECs, H1 cells were individualized with Accutase (Life Technologies), seeded in E8 with BMP4 (5ng/ml), Activin A (25ng/ml) and CHIR99021 (1 *μ*M) for the first 2 days, then withdraw CHIR99021 for the remaining period of differentiation. DECs were harvested at the end of day 5, sorting for CXCR4+ population for scRNA-seq experiments. For NPCs, the undifferentiated H1-SOX2-mCherry reporter line was treated with 0.5mM EDTA in PBS for 3 to 5 min and seeded in E6 (E8 minus FGF2, minus TGF*β*1), with 2.5 *μ*g/ml insulin, SB431542 (10 *μ*M) and 100 ng/ml Noggin. NPCs were harvested and enriched at the end of day 7, from sorting for the Cherry+ population for scRNA-seq experiments. All differentiation media were changed daily.

### Read mapping, quality control, and normalization

For each of the cell types studied, expected counts were obtained from RSEM [51]. In each condition there are a maximum of 96 cells, but all have fewer than 96 cells due to removal by quality control standards. Some cells were removed due to cell death or doublet cell capture, indicated by a post cell capture image analysis as well as a very low percentage of mapped reads. For more details on read mapping and quality control, see [47]. DESeq normalization [52] was carried out using the MedianNorm function in the EBSeq R package [53] to obtain library sizes. The library sizes were applied to scale the count data. Further, genes with very low detection rate (detected in fewer than 25% of cells in either condition) are not considered.

### Publicly available scRNA-seq datasets

Processed FPKM-normalized data from human myoblast cells [30] was obtained from GEO [54] using accession number GSE52529. In this study, we examined the set of cells cultured on standard growth medium (samples labeled with ‘T0’) as well as those treated with differentiation-inducing medium for 72 hours (samples labeled with ‘T72’). Processed TPM-normalized data from mESCs [31] was also obtained from GEO under accession number GSE60749. In this study, we examined the samples labeled as ‘mESC’ (cultured in standard medium), along with the samples labeled as ‘TwoiLIF’ (cultured in 2i+LIF differentiation-inhibitory medium).

### Publicly available bulk RNA-seq datasets

Modality of the gene expression distributions in bulk RNA-seq was explored using large, publicly available datasets and the results are displayed in Figure 2. In this figure, the red bars depict the bulk RNA-seq results, and datasets are labeled according to their source and sample size. Datasets GE.50, GE.75, and GE.100 are constructed by randomly sampling 50, 75, and 100 samples from GEUVADIS [55] in order to obtain sample sizes comparable to the single-cell sets under study (obtained from the GEUVADIS consortium data browser at www.ebi.ac.uk/arrayexpress/files/E-GEUV-1/analysis_results/GD660.GeneQuantCount.txt). Dataset LC consists of 77 normal lung tissue samples from the TCGA lung adenocarcinoma study [56] (obtained from GEO [54] using accession number GSE40419). All datasets were normalized using DESeq normalization [52] except for LC, for which the authors supplied values already normalized by RPKM.

### Mixture model formulation

#### Dirichlet Process Mixture of normals

Let 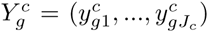 be the log-transformed nonzero expression measurements of gene *g* for a collection of *J*_*c*_ cells in condition *c* out of 2 total conditions. For simplicity of presentation, we drop the dependency on *g* for now, and let the total number of cells with nonzero measurements be *J*. We assume that under the null hypothesis of equivalent distributions (i.e. no dependency on condition), *Y* = {*Y*^*c*^}_*c*=1,2_ can be modeled by a conjugate Dirichlet process mixture (DPM) of normals given by

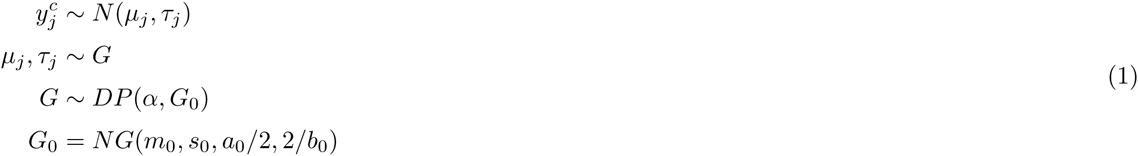

where *DP* is the Dirichlet process with base distribution *G*_0_ and precision parameter *α*, *N*(*μ*_*j*_, *τ*_*j*_) is the normal distribution parameterized with mean *μ*_*j*_ and precision *τ*_*j*_ (i.e. with variance 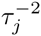), and *NG*(*m*_0_, *s*_0_, *a*_0_/2,2/*b*_0_) is the normal-gamma distribution with mean *m*_0_, precision *s*_0_*τ*, shape *a*_0_/2, and scale 2/*b*_0_. Let *K* denote the number of components (unique values among 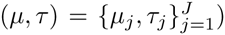. Note that two observations indexed by *j* and *j*′ are from the same cluster if and only if (*μ*_*j*_,*τ*_*j*_)=(*μ*_*j′*_,*τ*_*j′*_).

#### Product Partition Models

The posterior distribution of (*μ*,*τ*) is intractable even for moderate sample sizes. This is because the number of possible partitions (clusterings) of the data grows extremely rapidly as the sample size increases (according to the Bell number). However, if we let *Z* = (*z*_1_,…,*z*_*J*_) be the vector of component memberships of gene *g* for all samples, where the number of unique *Z* values is *K*, the probability density of *Y* conditional on *Z* can be viewed as a product partition model [57, 58]. Thus it can be written as a product over all cluster-specific component densities:

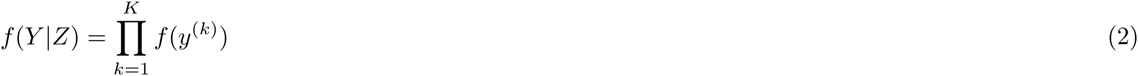

where *y*^(*k*)^ is the vector of observations belonging to component *k* and *f*(*y*^(*k*)^) is the component-specific distribution after integrating over all other parameters. In the conjugate normal-gamma setting, this has a closed form given by

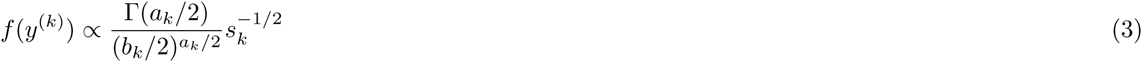

The posterior for the parameters (*μ*_*k*_, *τ*_*k*_) conditional on the partition is

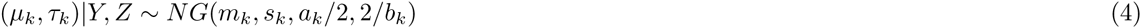

The posterior parameters (*m*_*k*_, *s*_*k*_, *a*_*k*_, *b*_*k*_) also have closed form due to the conjugacy of the model given by Equation 1. These parameters are given by

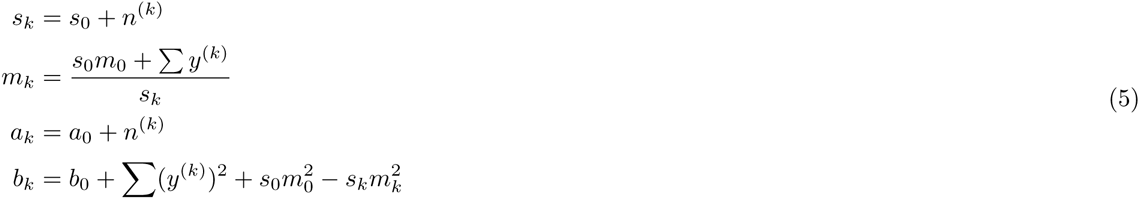

where *n*^(*k*)^ is the number of observations in cluster *k.* It follows that the marginal posterior distribution of *μ*_*k*_ conditional on the partition is

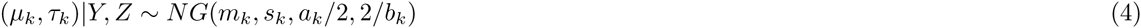

where *t*_*a*_(*b*, *c*) denotes the generalized Student’s t-distribution with *a* degrees of freedom, noncentrality parameter *b* and scale parameter *c*. The product partition Dirichlet process mixture model can be simplified as follows

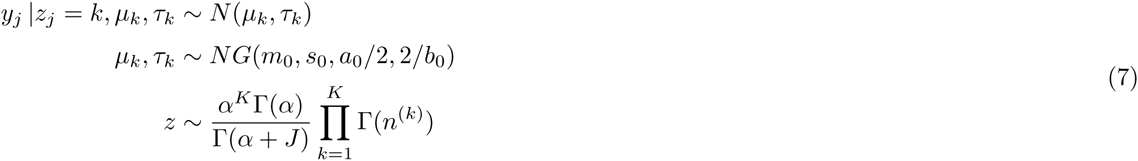

Then we can obtain the joint predictive distribution of the data *Y* and clustering *Z* by incorporating Equation 7:

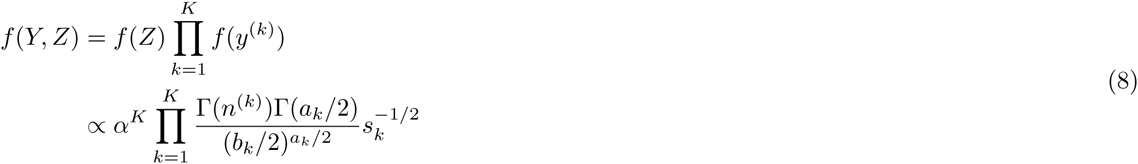

#### Model-fitting

The fitting of the model given in Equation Equation 7 involves obtaining an estimate *Ẑ* of the partition. The goal is to find the partition that yields highest posterior mass in Equation 8, referred to as the maximum a posteriori (MAP) partition estimate. Under this modeling framework, the solution for the MAP estimate is not deterministic and several computational procedures have been developed utilizing Polya urn Gibbs sampling [59, 60, 61], agglomerative greedy search algorithms [62, 63], or iterative stochastic search [64].

These procedures generally involve evaluation of the posterior at many different candidate partitions, and as such tend to be computationally intensive. To avoid this challenge, we recognize the relation to the corresponding estimation problem in the finite mixture model framework, where the partition estimate can be obtained by optimizing the BIC of the marginal density *f* (*Y*|*Z*) [65]. In fact, for certain settings of the prior distribution over partitions, the MAP estimate is identical to the estimate obtained by optimizing the BIC [58]. In practice, even when these settings are not invoked, the performance of partition estimates obtained via BIC optimization show comparable performance (see Supplement Section 1). We obtain the partition estimate *Ẑ* that optimizes the BIC using the `Mclust` R package [65] and satisfies the criteria for multi-modality described in the next section.

The hyperparameters for the cluster-specific mean and precision parameters were chosen so as to encode a heavy-tailed distribution over the parameters. Specifically, the parameters were set to *μ*_0_ = 0, 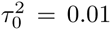, *a*_0_ = 0.01, and *b*_0_ = 0.01. The Dirichlet concentration parameter was set to *α* = 0.01, a choice of which is shown in Supplement Section 1 to be robust to many different settings in a sensitivity analysis.

#### Partition estimation

The partition estimate *Ẑ* is obtained that optimizes BIC using Mclust [65], in addition to the following filtering criteria. Note that the only constraint imposed on the number of components *K* in the modeling framework is that *K* ≤ *J*. However, under the sample sizes in this study, we only consider *K* ≤ 5. The first filtering criteria is based on the notion that a two-component mixture model is not necessarily bimodal [66], and relaxes the requirement that the MAP estimate correspond to the model with the lowest BIC. Specifically, for each candidate model fit by the BIC criterion with *K* components, a split step (if *K* = 1, obtain a new partition estimate *Z* with *K* = 2 unique elements) or a merge step (if *K* ≥ 2, obtain a new partition estimate *Ẑ* restricted to *K* – 1 unique elements) is carried out to generate a new candidate partition. The candidate partition with the larger value of *K* becomes the partition estimate only if the cluster separation suggests multi-modality. Cluster separation between any pair of clusters is assessed with the Bimodality Index (BI) [67]:

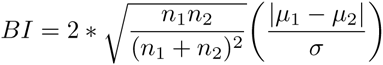

where the cluster means *μ*_1_ and *μ*_2_ are estimated via maximum likelihood, the common cluster standard deviation *σ* is conservatively estimated with the maximum cluster standard deviation among all clusters, and *n*_1_ and *n*_2_ are the number of cells belonging to each cluster. BI thresholds for the split and merge step were determined empirically and vary by sample size, as multiple modes are more easily detected as sample size increases [67] (for more details see Supplement Section 4).

The second filtering criteria is designed to reduce the impact of outlier cells. Specifically, clusters with fewer than 3 cells are not considered, and the merge step is also carried out if one of the clusters present has an extremely small variance (more than 20 times larger than any other cluster). Likewise, the split step is not carried out if one of the proposed clusters has a variance more than 10 times larger than any other cluster.

### Simulation details

#### Cluster means and variances

Each gene was simulated based on the characteristics of a randomly sampled unimodal gene with at least 25% nonzero measurements in the H1 dataset. For unimodal genes, the mean and variance were chosen to match the observed mean and variance; for bimodal genes, the cluster means and variances were selected to be near the observed mean and variance. The proportion of zeroes is chosen to match that observed in the randomly sampled gene, and is not varied by condition. Details are provided in the following sections.

Distances between (log-scale) component means Δ_*μ*_*σ̂* in the multi-modal genes were chosen such that clusters were separated by a minimum of 2 and maximum of 6 standard deviations, where the standard deviation *σ* is assumed constant (on the log-scale) across clusters and is estimated empirically assuming a lognormal distribution on the raw scale. In this setting, the cluster distance can also be thought of as a fold-change within condition (across clusters), where the ratio of the cluster means (untransformed-scale) is equal to *e*^Δ^_*μ*_*^σ̂^*. The ratio of the cluster standard deviations (raw-scale) is also equal to this same fold change (see Supplement Section 2.1 for more details). The cluster mean distance values were chosen to represent a range of settings for which the difficulty of detecting multi-modality is widely varied, as well as to reflect the range of observed cluster mean distances detected empirically in the case studies.

#### Unimodal genes

Parameters of the negative binomial distribution for unimodal genes were estimated from the randomly sampled observed genes using the method-of-moments. These empirical parameters were used as is to simulate both conditions of EE genes, and condition 1 of DE and DB. Condition 1 of DM was simulated by decreasing the mean by half the value of Δ_*μ*_. The second condition for DE genes was simulated based on condition 1 parameters using randomly sampled fold changes that were between 2 and 3 standard deviations of the observed fold changes between H1 and DEC.

#### Bimodal genes

Parameters for the mixture of negative binomial distributions in bimodal genes were also generated using empirically estimated means and variances. The first (lower) component mean was decreased by half the value of Δ_*μ*_ and the second (higher) component mean was increased by half the value of Δ_*μ*_.

### DD classification algorithm

Genes detected as significantly DD from the permutation test of the Bayes Factor score are categorized into patterns of interest. The genes that are not classified as either DE, DP, DM, or DB are considered ‘no calls’, abbreviated NC. These represent patterns that are not of primary interest, such as those with the same number of components within each condition and overall, but not significantly different cluster-specific means. Genes with this pattern that are significantly DD could arise if, for example, the cluster-specific variances differ across conditions. We do not infer differences of these types since it is possible that they could be explained by cell-specific differences in technical variation [17].

An additional step to improve the power to detect genes in the DP category was also implemented. This step was motivated by the observation that the Bayes Factor score tends to be small when the clustering process within each condition is consistent with that overall, as in the case of DP. Thus, for genes that were not significantly DD by permutation but had the same number of components within condition as overall, Fisher’s exact test was used to test for independence with biological condition. If the p-value for that test is less than 0.05, then the gene was added to the DP category (this did not result in the addition of any false positives in the simulation study). In addition, since the Bayes Factor score depends on the estimated partition, we increase the robustness of the approach to detect DD genes under possible misspecification of the partition by also assessing evidence of DD in the form of an overall mean shift for genes not significant by the permutation test (using a t-statistic with FDR controlled by [34]). This resulted in the detection of between 121 and 689 additional genes in the hESC comparisons and did not add any false positives in 94% of simulation replications (with only a single false positive gene in the other 6% of replications).

Here we present pseudocode for the classification of DD genes into the categories DE, DP, DM, or DB. For every pair of clusters, obtain a sample of 10,000 observations from the posterior distribution of the difference in means. The clusters are considered to overlap if the 100% credible interval contains 0.

#### DD classification algorithm

**if c**_**1**_ = **c**_**2**_ = **1**

**if** clusters *c*_1_ and *c*_2_ do not overlap ⇒ DE

**else** ⇒ NC

**else if c**_**1**_ = *c*_**2**_ ≥ **2**

**if c**_1_ = **c**_2_ = **c**_**OA**_

**if** At least *c*_1_ of the clusters overlap ⇒ DP

**else** ⇒ NC

**else if c**_1_ = **c**_2_ < **c**_OA_

**if** at most one cluster pair overlaps ⇒ DE

**else** ⇒ NC

**else if c**_**1**_ ≠ **c**_**2**_

**if** no cluster pairs overlap ⇒ DB

**else** ⇒ DM

## Availability of supporting data

The hESC data has been deposited in GEO [54] with accession number GSE75748.

Sensitivity analyses, further methodological details, and additional results are provided in a supplement.

## List of abbreviations

scRNA-seq: single-cell RNA sequencing
scDD: single-cell differential distributions DE: differential expression
DP: differential proportion
DM: differential modality
DB: differential both (expression and modality)
DZ: differential zeroes
hESC: human embryonic stem cell
DEC: definitive endoderm cell
NPC: neuronal progenitor cell
DPM: Dirichlet process mixture
PPM: product partition model
MAP: maximum a posteriori
BIC: Bayesian information criterion

## Competing interests

The authors declare that they have no competing interests.

## Author’s contributions

CK, L-FC, JAT, RMS, and KDK formulated the problem. CK, KDK, and MAN developed the scDD model. KDK implemented the scDD model in R, developed and implemented simulations, and applied scDD to the hESC case study data. YL assisted with the simulation study. L-FC conducted the hESC experiments. CK, L-FC, RMS, and KDK interpreted results. KDK and CK wrote the paper. All authors read and approved the final manuscript.

## Acknowledgements

This work was supported by NIH GM102756 (CK), NIH U54AI117924 (CK), NIH 4UH3TR000506-03 (JAT), and 5U01HL099773-06 (JAT). The authors thank the editorial staff and two anonymous reviewers for insightful comments and suggestions that helped improve the quality of the manuscript.

## Additional Files

Additional file 1 — Supplement

Sensitivity analyses of MAP estimation method, further methodological details, and additional results.

